# PgRC: Pseudogenome based Read Compressor

**DOI:** 10.1101/710822

**Authors:** Tomasz Kowalski, Szymon Grabowski

## Abstract

**Motivation:** The amount of sequencing data from High-Throughput Sequencing technologies grows at a pace exceeding the one predicted by Moore’s law. One of the basic requirements is to efficiently store and transmit such huge collections of data. Despite significant interest in designing FASTQ compressors, they are still imperfect in terms of compression ratio or decompression resources.

**Results:** We present Pseudogenome-based Read Compressor (PgRC), an in-memory algorithm for compressing the DNA stream, based on the idea of building an approximation of the shortest common superstring over high-quality reads. Experiments show that PgRC wins in compression ratio over its main competitors, SPRING and Minicom, by up to 18 and 21 percent on average, respectively, while being at least comparably fast in decompression.

**Availability:** PgRC can be downloaded from https://github.com/kowallus/PgRC.

## 1 Introduction

High-throughput sequencing, producing hundreds of millions of short reads in a typical experiment, is still the dominating technology on the market, despite a notable progress in long-read sequencing. Major repositories nowadays store petabases of DNA and RNA sequence information (e.g., over 29 petabases in NIH Sequence Read Archive as of July 2019) and grow rapidly.

Raw sequencing data are typically stored in FASTQ format as a sequence of records (reads). Each read basically consists of three textual fields: its identifier (header), a DNA string and the corresponding quality score string; the last two sequences are of the same length. To maintain the enormous amounts of FASTQ data, efficient data compression techniques must be utilized.

The early dedicated solutions, proposed almost a decade ago (Tembe *et al*., 2010; Deorowicz and Grabowski, 2011; Bonfield and Mahoney, 2013), were better than general-purpose compressors, like gzip, but not dramatically so. Parallelization (Howison, 2013; Roguski and Deorowicz, 2014) made the compression techniques more practical, but the compression ratio was still hampered with the quasi-random nature of quality scores and the difficulty of finding long matches between DNA strings in large datasets. In particular, using the listed methods the DNA symbols could not be compressed much better than with the naive 2 bits per base. Two observations were the basis of breakthroughs that followed. According to one, the quality data had only minor role in further analyses, like variant calling, and for this reason lossy compression could be successfully applied to this stream (Ochoa *et al*., 2017; Roguski *et al*., 2018). Note that in accordance with those discoveries (more references can be found in (Ochoa *et al*., 2017)), the output of recent Illumina sequencers typically contains quality scores binned (quantized) to 8 levels only.

The other general observation was that non-standard similarity detection techniques applied to the DNA stream can eliminate its redundancy due to overlapping reads in a significant degree. Those non-standard techniques, elaborated in the following paragraphs, are either disk-based or memory-based. We focus on the compression of the DNA stream (some of the algorithms described below are limited to those data).

ReCoil (Yanovsky, 2011) creates a similarity graph for the input dataset, defined as a weighted undirected graph with vertices corresponding to the reads of the dataset. The edge weight between a pair of reads, *R*_1_ and *R*_2_, is related to the “profitability” of storing *R*_1_ and the edit script transforming *R*_1_ into *R*_2_ versus storing both reads explicitly. Then, a minimum spanning tree (MST) procedure is run, during which each node is encoded using the set of maximum exact matches between the node’s read and the read of its parent in the MST. The ReCoil approach explored “new” redundancies with a progress in compression ratio, but unfortunately, was computationally costly and thus hardly scalable. BEETL (Cox *et al*., 2012) employs the Burrows–Wheeler Transform (BWT) to group similar reads, which are then compressed with general-purpose compression techniques. Interestingly, both ReCoil and BEETL—despite widely different underlying ideas—made use of disk space and thus were not limited by the amount of available RAM memory.

SCALCE (Hach *et al*., 2012) partitions the reads into buckets, based on a combinatorial pattern matching technique called locally consistent parsing. Reads with a large overlap are likely to have the same core string and thus are sent to the same bucket. The contents of each bucket are then gzip-compressed.

ORCOM (Grabowski *et al*., 2015) borrows some ideas from its predecessors. It is a disk-based algorithm which reorders the input reads and distributes them into buckets. Its key concept, however, is to use so-called minimizers (Roberts *et al*., 2004) for the bucket labels. A minimizer of length *ℓ* for a read *R* of length *m* is the lexicographically smallest of the (*m* − *ℓ* + 1) *ℓ*-mers of *R*. A canonical minimizer, which is actually used by ORCOM, is a minimizer taken over the read and its reversed-complemented form. Two reads with a large overlap are likely to share the same (canonical or non-canonical) minimizer and thus the same bucket. The contents of each bucket are compressed separately, with sorting the reads from their minimizer’s position, careful modeling of mismatches and other minor improvements, combined with arithmetic coding or PPMd (context-based) compression applied to several resulting data streams. The compression ratio on a 134 Gbp human genome sequencing data achieved by ORCOM was 0.317 bits per base, improving the BEETL’s result of 0.518 bits per base.

Mince (Patro and Kingsford, 2015) is a related algorithm, but its distribution of reads into buckets is based on the number of shared *k*-mers. More precisely, a read *R* is thrown to the bucket which maximizes the number of *k*-mers of *R* occurring in any read the bucket contains. Its compression ratio is in most cases by a few percent higher than ORCOM’s (see, e.g., extensive comparisons in (Liu *et al*., 2018)), but is less efficient in terms of time and memory usage. FaStore (Roguski *et al*., 2018) also follows the ORCOM approach, but improves its compression ratio (by a factor of about 1.2 typically) mostly thanks to re-distribution of reads from the buckets and assembling reads into contigs; in other words, it allows to merge similar clusters of reads. FaStore also boasts with good performance—decompression speed exceeding 100 MB/s, and even 250 MB/s in one of the modes, using 8 threads—and several lossy modes for the quality and header streams.

HARC (Chandak *et al*., 2018a) resigns from disk-based bucketing, in favor of a succinct in-memory hash tables. Its basic idea is to find maximum overlaps between reads and create consensus sequences, using majority voting over mismatching nucleotides in groups of reads. HARC’s compression ratios are typically higher than FaStore’s. SPRING (Chandak *et al*., 2018b), from (mostly) the same authors, is a follow-up tool, again with some (relatively small) improvements in compression of the DNA stream, and also capable to handle all FASTQ streams.

Minicom (Liu *et al*., 2018) takes an approach similar to FaStore, but is more effective in producing contigs. It creates contigs for groups of reads having the same minimizer and then merges them across groups based on a suffix-prefix overlap index. It works with relatively long minimizers, e.g., 31 symbols.

FQSqueezer (Deorowicz, 2019), presented very recently, beats SPRING and Minicom by even tens of percent in compression ratio, but is impractical, especially due to decompression times up to two orders of magnitude longer than of its main competitors, and large memory requirements. Its PPM-based approach, with a dedicated error correction technique, shows however the current limit of read compression.

It has been early recognized that an algorithm can achieve optimal compression if reads are reordered according to their position in the original genome. Indeed, many of the algorithms listed above search for overlaps, and then possibly long contigs, among the reads. Yet, there exists a group of FASTQ compressors which attempt de novo assembly more explicitly. A challenging phase of such algorithms is to construct de Bruijn graphs. A progress in this vein, with probabilistic de Bruijn graphs construction (based on the Bloom filter data structure), was witnessed by Quip (Jones *et al*., 2012) and LEON (Benoit *et al*., 2015). Even better compression (but still not on a par with, e.g., SPRING and Minicom) was achieved in Assembltrie (Ginart *et al*., 2018), where light de novo assembly was essentially performed via storing reads in a compact trie.

So far we discussed reference-free compression methods. Another line of research assumes that a reference genome is available, which essentially allows to replace each read with a pointer to the reference genome, together with an edit script marking the discrepancies between the read and its locus. There are several compressors following this path, including (Fritz *et al*., 2011; Zhang *et al*., 2015); let us briefly present only two non-standard ones. PathEnc (Kingsford and Patro, 2015) does not align reads to the reference (which is a relatively costly process); instead, it only uses the reference as the model during the (context-based) compression of reads. The model is constantly updated during the (de)compression with the given data, so even a poorly adequate reference sequence may bring about some compression gain. Quark (Sarkar and Patro, 2017), designed for RNA-seq, uses the reference for the compression only. On the decompression side, the reference sequence is not needed, as everything needed to decompress the reads is stored in the final compressed file, as a collection of small subsets (“islands”) of the transcriptome. It would seem that lack of a reference sequence should seriously handicap reference-free compressors, in comparison. Surprisingly though, PathEnc and Quark, which belong to the best reference-based compressors, only rarely produce smaller archives than top reference-free tools (Minicom, SPRING, HARC), as demonstrated in (Liu *et al*., 2018). This may suggest that the area of reference-based compressors is actually underdeveloped.

In this work we propose a new reference-free compressor for the DNA stream of FASTQ files, called Pseudogenome-based Read Compressor (PgRC), which also exploits read overlaps. Its attained compression ratios surpass SPRING and Minicom, preserving fast decompression.

## 2 Methods

### 2.1 Overview

We assume that the input reads are of fixed length, over the alphabet `ACGTN`. The description in this and following subsections concerns the most basic compression mode, SE, which means that the input data are single-end (SE) reads and their given order will not be preserved. The other compression modes (SE_ORD, PE, PE_ORD) share most of the concepts of the basic scheme and the modifications are presented in Sect. 2.6.

The proposed PgRC is based on a few ideas. The key one is an approximation of the shortest common superstring over a set of the given reads, which we call a “pseudogenome” (hence the name of our tool), an idea basically described in (Kowalski *et al*., 2015). In this work, however, we modify the procedure from our earlier research, by partitioning the read set into groups, related to their quality and the existence of `N` symbols in them.

The high-quality pseudogenome, let it be denoted by *PG*_*hq*_, is built from the reads in their forward orientation. Note that knowing the pseudogenome and the mapping locations (offsets) of the reads is enough to reconstruct the (high quality) reads. The low quality reads, i.e., those which did not participate in the *PG*_*hq*_ construction, are (tried to be) mapped to *PG*_*hq*_ as well, but in a relaxed way. Firstly, approximate matching is performed with several allowed mismatches (Hamming errors) between a read and the corresponding pseudogenome’s area. Secondly, the low quality reads are tried to be mapped twice: directly and in the reverse-complemented form. From those low quality reads which do not admit successful mapping we construct another pseudogenome (*PG*_*lq*_).

It appears that both *PG*_*hq*_ and *PG*_*lq*_ have internal redundancy, as well as matching strings between one and the other of these sequences. Moreover, significant part of this redundancy consists of reverse-complemented repetitions. We denote reverse-complemented sequence *S* by *S*^*rc*^. As direct matches will be handled in a further processing phase (by a general-purpose compressor), we focus on reverse-complemented matches only. To this end, we find and encode matches between *PG*_*hq*_ and 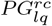, and between *PG*_*hq*_ and 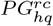.

As a result of the sketched processing, several data components are produced, which are then compressed using a general-purpose compressor. The following subsections present details of the algorithm.

### 2.2 Read partitioning and pseudogenome generation

As mentioned in the overview, reads should be partitioned into those of high and of low quality. To this end, two ideas were considered. One was to estimate the quality of each read based on its quality scores, as simply the product of occurrence probabilities for all its bases, followed by filtering out “bad” ones using a specified threshold. Although this idea requires side information for encoding the DNA stream, in practice it poses no problem as the input is full FASTQ data. Further experiments (see the Supplementary data) showed however that even better compression ratios can be obtained with different means.

More concretely, we followed the pseudogenome construction algorithm (Kowalski *et al*., 2015). Given a read array 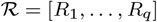, where *R*_*i*_ = *R*_*i*_[1 … *m*] for all *i*, we define a pseudogenome as a sequence *PG*[1 … *g*] such that for all *i* there is such *j*_*i*_ that *R*_*i*_ = *PG*[*j*_*i*_ … *j*_*i*_ + *m* − 1]. Obviously, we try to minimize *g*. Finding the shortest pseudogenome in string matching literature is call the shortest common superstring (SCS) problem and is known to be NP-hard (Maier and Storer, 1997). Our heuristic solution obtains a pseudogenome in *O*(*qm*(*m* + log *q*)) time, with a possibility of removing the logarithmic factor when quick sort is replaced with radix sort. The algorithm produces read overlaps, starting from the longest ones. The modification in the current work, for the purpose of read partitioning, is to terminate the procedure once the length of the found overlaps reaches a specified threshold *ov*, 65% of the read length by default. In another pass over the read collection we keep only those reads for which we previously found (long enough) *both left and right* overlaps.

The remaining reads do not participate in *PG*_*hq*_ generation. In particular, those reads which contain at least one symbol N are separated out once they are read from the dataset, which is beneficial both for compression speed and memory usage.

### 2.3 Read alignment

In this phase we try to align those reads which have not participated in *PG*_*hq*_ construction onto it. Most of those matches will be approximate, with mismatches. We set the maximum error level, i.e., the maximum allowed number of mismatches per aligned read, to ⌊*read_len/M*⌋, where *M* = 6 by default (which means, for example, that for reads of length 100 we allow up to 16 mismatches). Each successfully aligned read will be represented as an offset (position) in *PG*_*hq*_ at which the aligned read starts, the number of mismatches and their relative locations, the read’s symbols at the mismatching positions, as well as a flag telling if the read is mapped forwardly or reverse-complemented. Mismatch locations are sent to multiple (conceptual) streams, according to the number of mismatches, i.e., locations for reads with one mismatch are dispatched to the first stream, locations for reads with two mismatches to the second stream, etc. Additionally, the mismatch locations are given in the reversed order (to exploit the asymmetric error distribution in reads), with the mismatch exactly at the last symbol denoted as location 0, and differentially encoded. As two different mismatches cannot have the same location, the differences between successive mismatches are decremented by 1. To illustrate, if a read of length 100 is mapped with 3 mismatches, at the positions: 70, 75 and 100, the positions are encoded as: 0 (= 100 − 100), 24 (= 100 − 75 − 1), 4 (= 75 − 70 − 1). Another example is given in Fig. 1. Note that different streams have somewhat different distributions, as the average values (gaps) in, e.g., tenth mismatch stream are smaller than in the second stream.

**Fig. 1.**
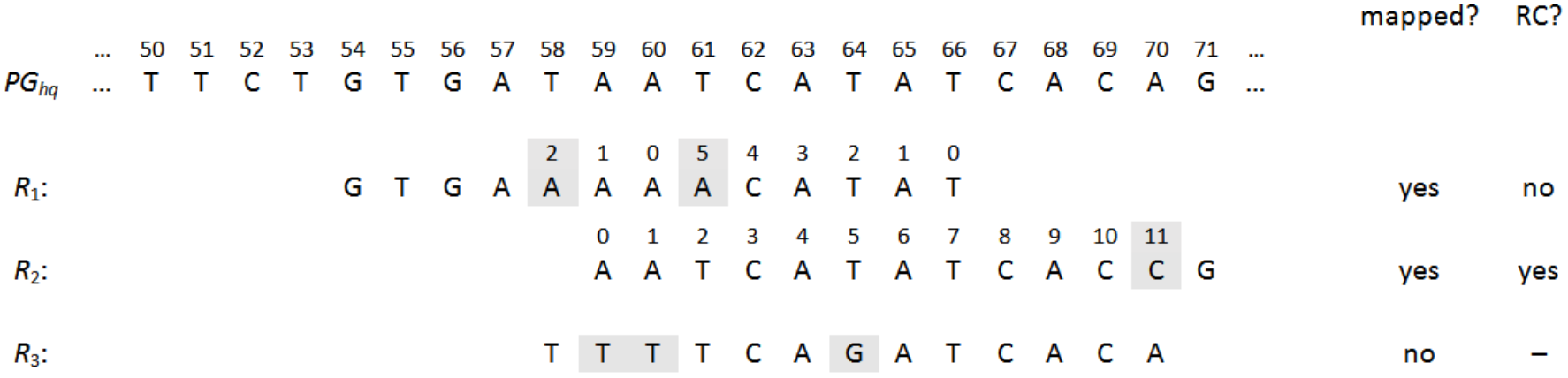
Example of aligning three reads, *R*_1_ = GTGAAAAACATAT, *R*_2_ = CGGTGATATGATT, and *R*_3_ = TTTTCAGATCACA, onto the reference sequence *PG*_*hq*_. The reads are of length 13; we set *M* = 6, i.e., we allow up to ⌊13/6⌋ = 2 mismatches. *R*_1_ is aligned in forward, while *R*_2_ is reverse-complemented, as stated in the rightmost column. *R*_3_ cannot be aligned within the error limit. The mismatching characters in reads are marked. Their differentially encoded positions are marked above the corresponding symbols.

To find alignments efficiently, we build a hash table with *k*-mers sampled from *PG*_*hq*_ and for each candidate read we find a seed of length *s*, where *s* ≥ *k* (by default, *s* = 38 and *k* = 28), such that *s* successive symbols from the read match exactly some area in *PG*_*hq*_. This is performed efficiently using the sparse sampling technique from copMEM (Grabowski and Bieniecki, 2019), a recently proposed tool for maximum exact match finding. A reasonable time-compression tradeoff in this phase is obtained by setting the maximum allowed number of items inserted per hash (it was experimentally set to 12); further *k*-mers hashing to a full slot are simply not inserted. The number of collisions in the hash table is thus in many cases significantly reduced with a negligible loss in compression.

### 2.4 Squeezing the pseudogenomes

From the low-quality reads which have not mapped onto *PG*_*hq*_ we create two pseudogenomes, *PG*_*lq*_ and *PG*_*N*_, where the latter is based on reads containing symbols `N`. The high-quality pseudogenome, *PG*_*hq*_, has specific redundancy, which cannot be removed by general-purpose compressing techniques (e.g., the LZMA algorithm). *PG*_*hq*_ contains reverse-complemented matches of quite long strings. To give an example, the string `AAA…C` of length 60 (59 symbols `A` followed by one `C`), with a single occurrence in *PG*_*hq*_, could be removed, provided that `GTTT…T` (one `G` followed by 59 symbols `T`) occurs earlier in *PG*_*hq*_. Obviously, the positions and lengths of the removed strings, as well as the positions of the matched strings, must be marked in a way to make reconstruction of the pseudogenome possible. The existence of the reverse-complemented repetitions can be explained by the nature of the input reads, which come from both DNA strands. Moreover, similar redundancy exists between *PG*_*hq*_ and *PG*_*lq*_.

To remove the repetitions, with a noticeable improvement in compression, we again use the copMEM sparse sampling technique. Reverse-complemented repeats of length at least *p* (where *p* was set to 50 by default) are found using a hash table. The copies are removed from a given pseudogenome and their representations as two values, match offset and match length (the latter decreased by *p*), are sent to separate streams.

### 2.5 Compression of the by-products

The by-products obtained in the aforedescribed stages still contain some redundancy. In case of the three pseudogenomes, *PG*_*hq*_, *PG*_*lq*_ and *PG*_*N*_, there exist both LZ77 redundancy and statistical redundancy. The former means that there occur repeating sequences (matches) in a concatenation of the three pseudogenomes, which essentially can be encoded as pairs (offset, length). The latter is addressed with compact encoding of symbols (literals), according to the rule that shorter codewords should be assigned to frequent symbols, and longer codewords to the rarer ones, and can be achieved, e.g., with Huffman coding. The compression algorithm we chose for squeezing the pseudogenomes is LZMA, incorporated in the popular 7zip tool. LZMA combines strong LZ77 with statistical compression and is capable of finding matches in a sliding window of size up to 1.5 GB.

LZMA is used also to compress match offsets and lengths of the reverse-complemented repeats, where the input lengths are variable-length and take either 1 byte if the length, decreased by the threshold *p*, is at most 127, or 2 bytes if the length minus *p* is at most 2^14^ − 1 = 16383, and so on. In practice, in most cases the length needs 1 byte.

The remaining by-products do not yield significant LZ77 redundancy, but it appears that they have some contextual redundancy, which can be effectively removed with a low-order PPM compressor. To this end, we applied PPMd order-2 for them, again from the 7zip tool.

A couple of words are needed to describe their performance characteristics. LZMA is relatively slow at compression, especially using a large sliding window and compression-boosting options. On the other hand, it is rather fast in decompression, reaching the speed of at least tens of megabytes per second. PPMd is a symmetric algorithm, which means that both compression and decompression work at about the same speed, often attaining more than 10 MB/s. The PPMd speed worsens with higher order, but in our usage scenario order-2 seemed to be the best choice (order-3 and higher do not help in compression, or help very slightly, but are more computationally demanding).

LZMA and PPMd in 7zip are one-threaded, both in compression and decompression.

### 2.6 Other modes

As stated, the previous subsections describe handling the SE reads without preserving their order. Now we focus on the other compression modes. If the order of reads is not to be preserved (i.e., in SE and PE modes), then the reads are written in the order of their occurrence position in a respective pseudogenome (*PG*_*hq*_, *PG*_*lq*_, or *PG*_*N*_), and those positions are differentially encoded. If a given read is part of *PG*_*hq*_, in the same order its satellite data are stored: read orientation and mismatch information. (Note that reads belonging to *PG*_*lq*_ to *PG*_*N*_ cannot be reverse-complemented or aligned with mismatches.)

In the PE mode we need to store the pairing information. Like other tools, PgRC also keeps track which of the reads of a pair is taken from the first (_1) and which from the second (_2) PE file.

In the modes PE and PE_ORD, the reads from the file _2 are reverse-complemented at the start. Thanks to it, in most cases for high-quality reads both reads of the pair will be located close to each other in *PG*_*hq*_. This little trick increases the chance for a good mapping for both reads, which is clearly beneficial for efficient encoding of the pairing information, and may also make the pseudogenome more compact. The pairing information is split into as many as 7 streams for careful encoding (see details in the Supplementary data).

In the order preserving modes, SE_ORD and PE_ORD, reads are sent to the output in their original order (disregarding their order in a pseudogenome). In the SE_ORD mode all these positions are stored in a single stream, compressed with LZMA. In PE_ORD an analogous stream is maintained and LZMA-compressed only for the reads from the first file, the information about their mates in the second file is decoded from pairing information.

## 3 Results

PgRC was tested on a Linux machine equipped with an 6-core Intel Core i7-7800X 3.5 GHz CPU, 128 GB of DDR4-RAM (CL 16, clocked at 2666 MHz) and a 5 TB HDD (Toshiba X300 HDWE150EZSTA).

The test collection consists of a number of real genome and transcriptome sequencing datasets, used for benchmarking in prior works on FASTQ compression. Their URLs and details can be found in the Supplementary data. All, original and compressed, dataset sizes are given in GBytes (or Gigabases), where *G* = 10^9^. SPRING and Minicom compressors were run with 12 threads (the test CPU sports 12 hardware threads, thanks to hyper-threading) and other settings default. PgRC is a single-threaded implementation.

Tables 1 and 2 present the results in the order non-preserving and order-preserving SE modes, respectively (SE and SE_ORD in short). The table rows are sorted from the largest to the smallest dataset. In both experiments PgRC wins in compression ratio over the stronger of its competitors in almost all cases, namely in 13 out of 15 in both SE and SE_ORD modes.

**Table 1.** Compression, time and memory usage in the order non-preserving regime on SE datasets

| dataset | size<br>[Gbs] | SPRING |  |  |  |  | Minicom |  |  |  |  | PgRC |  |  |  |  |
| --- | --- | --- | --- | --- | --- | --- | --- | --- | --- | --- | --- | --- | --- | --- | --- | --- |
|  |  | ratio | ctime | dtime | cmem | dmem | ratio | ctime | dtime | cmem | dmem | ratio | ctime | dtime | cmem | dmem |
| ERP001775_1 | 44.39 | 0.2283 | 6283.0 | 1800.8 | 23.48 | 5.28 | 0.3316* | 23553.0 | 1160.2 | 114.20 | 4.07 | 0.2182 | 17410.0 | 423.4 | 36.92 | 16.43 |
| ERR174310_1 | 20.97 | 0.4096 | 2513.3 | 217.8 | 11.27 | 4.93 | 0.5888* | 14778.0 | 222.0 | 68.80 | 4.07 | 0.4527 | 15197.0 | 135.5 | 21.17 | 12.35 |
| SRR870667_1 | 7.48 | 1.2924 | 1135.2 | 182.4 | 7.42 | 3.12 | 1.2117 | 2813.4 | 122.3 | 27.71 | 4.07 | 0.7636 | 14151.0 | 79.2 | 19.95 | 13.76 |
| SRR445724 | 5.09 | 0.4328 | 685.8 | 70.0 | 3.47 | 2.52 | 0.3860* | 1222.5 | 43.1 | 13.04 | 1.37 | 0.3444 | 2870.5 | 29.7 | 8.59 | 2.97 |
| SRR445726 | 4.92 | 0.3882 | 581.6 | 47.3 | 3.61 | 2.48 | 0.3507* | 1186.3 | 39.4 | 12.24 | 1.26 | 0.3132 | 2472.8 | 27.3 | 7.57 | 2.59 |
| SRR490961 | 4.91 | 0.2183 | 467.3 | 40.9 | 2.92 | 2.15 | 0.1995* | 847.5 | 27.3 | 11.02 | 0.97 | 0.1841 | 1343.3 | 22.6 | 3.77 | 1.29 |
| SRR1294116 | 4.64 | 0.2763 | 304.9 | 158.8 | 3.19 | 2.35 | 0.2695 | 411.1 | 38.1 | 10.54 | 0.69 | 0.2415 | 1495.8 | 28.3 | 4.44 | 1.45 |
| ERR532393_1 | 3.58 | 0.4325 | 169.3 | 35.8 | 3.04 | 1.85 | 0.4098 | 457.4 | 26.3 | 10.47 | 0.69 | 0.3765 | 1749.7 | 21.9 | 6.18 | 1.36 |
| SRR065390_1 | 3.38 | 0.2275 | 175.3 | 28.6 | 2.63 | 1.24 | 0.2565 | 516.5 | 24.3 | 8.82 | 0.64 | 0.1877 | 773.1 | 17.9 | 2.09 | 0.79 |
| SRR490976 | 3.33 | 0.3851 | 407.2 | 32.4 | 2.84 | 2.14 | 0.3527 | 421.4 | 30.7 | 8.45 | 0.69 | 0.3411 | 1684.9 | 20.5 | 5.33 | 1.84 |
| SRR445718 | 3.29 | 0.3371 | 315.8 | 31.7 | 2.65 | 2.03 | 0.3062* | 396.8 | 24.6 | 8.27 | 0.69 | 0.2855 | 1365.7 | 18.6 | 4.02 | 1.33 |
| SRR689233_1 | 1.48 | 0.1934 | 66.0 | 14.1 | 1.61 | 0.80 | 0.1874 | 106.4 | 11.7 | 4.59 | 0.29 | 0.1700 | 330.9 | 6.2 | 0.90 | 0.33 |
| SRR635193_1 | 1.47 | 0.2671 | 108.2 | 16.2 | 1.30 | 0.81 | 0.2887 | 396.9 | 13.0 | 5.62 | 0.35 | 0.2599 | 438.1 | 7.9 | 1.78 | 0.53 |
| MiSeq_1 | 0.87 | 0.1253 | 36.6 | 7.3 | 1.66 | 0.84 | 0.1108 | 84.0 | 3.5 | 2.70 | 0.15 | 0.0947 | 110.1 | 2.2 | 0.39 | 0.11 |
| SRR554369_1 | 0.17 | 0.2397 | 7.2 | 1.8 | 0.86 | 0.32 | 0.3073 | 23.9 | 1.2 | 1.74 | 0.03 | 0.2409 | 32.2 | 0.7 | 0.23 | 0.06 |

Compression ratios are in bits per base (bpb). “ctime” (resp. “dtime”) stands for compression (resp. decompression) time, and is expressed in seconds. “cmem” (resp. “dmem”) stands for peak compression (resp. decompression) memory usage and is expressed in gigabytes ( $=10^9$ bytes). Minicom failed to correctly decompress several datasets, which are marked with an asterisk. The name “MiSeq\_1” denotes the dataset MiSeq\_Ecoli\_MG1655\_110721\_PF\_1.

**Table 2.** Compression, time and memory usage in the order preserving regime on SE datasets

| dataset | size<br>[Gbs] | SPRING |  |  |  |  | Minicom |  |  |  |  | PgRC |  |  |  |  |
| --- | --- | --- | --- | --- | --- | --- | --- | --- | --- | --- | --- | --- | --- | --- | --- | --- |
|  |  | ratio | ctime | dtime | cmem | dmem | ratio | ctime | dtime | cmem | dmem | ratio | ctime | dtime | cmem | dmem |
| ERP001775_1 | 44.39 | 0.5275 | 4772.0 | 1426.8 | 23.48 | 4.30 | 0.6251* | 23672.0 | 717.5 | 114.26 | 52.74 | 0.5039 | 22530.0 | 369.8 | 40.44 | 19.51 |
| ERR174310_1 | 20.97 | 0.6949 | 2509.0 | 248.5 | 11.27 | 4.43 | 0.8573* | 14852.0 | 352.4 | 69.13 | 24.91 | 0.7236 | 17571.0 | 205.3 | 25.12 | 13.81 |
| SRR870667_1 | 7.48 | 1.4547 | 826.3 | 143.2 | 6.88 | 2.31 | 1.3638 | 2963.0 | 159.5 | 27.78 | 9.42 | 1.0332 | 14604.0 | 102.7 | 20.48 | 14.24 |
| SRR445724 | 5.09 | 0.6574 | 606.4 | 64.6 | 2.90 | 1.12 | 0.6074* | 1224.0 | 82.0 | 13.04 | 6.11 | 0.5984 | 2887.1 | 42.4 | 9.00 | 3.13 |
| SRR445726 | 4.92 | 0.6130 | 615.7 | 52.9 | 2.83 | 1.07 | 0.5721* | 1202.7 | 78.0 | 12.23 | 5.91 | 0.5646 | 2505.7 | 39.4 | 7.97 | 2.73 |
| SRR490961 | 4.91 | 0.4572 | 423.0 | 47.2 | 2.83 | 0.83 | 0.4398* | 855.5 | 74.2 | 11.01 | 5.90 | 0.4336 | 1411.5 | 43.5 | 4.17 | 1.44 |
| SRR1294116 | 4.64 | 0.5122 | 213.3 | 87.7 | 2.70 | 0.90 | 0.5022 | 456.1 | 72.8 | 10.55 | 5.52 | 0.4829 | 1579.1 | 42.7 | 4.80 | 1.59 |
| ERR532393_1 | 3.58 | 0.6667 | 166.7 | 40.1 | 1.78 | 1.00 | 0.6469 | 464.1 | 62.6 | 10.44 | 4.29 | 0.6206 | 1774.5 | 30.2 | 6.47 | 1.47 |
| SRR065390_1 | 3.38 | 0.4776 | 175.4 | 32.7 | 2.63 | 0.83 | 0.5050 | 560.7 | 56.5 | 8.74 | 4.06 | 0.4278 | 805.2 | 19.4 | 2.47 | 0.89 |
| SRR490976 | 3.33 | 0.5979 | 435.3 | 36.6 | 2.23 | 1.11 | 0.5665 | 420.6 | 55.2 | 8.42 | 4.00 | 0.5879 | 1730.5 | 28.7 | 5.60 | 1.94 |
| SRR445718 | 3.29 | 0.5704 | 229.6 | 35.4 | 2.21 | 0.89 | 0.5355 | 403.9 | 55.6 | 8.24 | 3.96 | 0.5346 | 1392.4 | 32.8 | 4.28 | 1.43 |
| SRR689233_1 | 1.48 | 0.4489 | 66.1 | 14.5 | 1.61 | 0.74 | 0.4419 | 109.4 | 20.9 | 4.59 | 1.97 | 0.4303 | 347.5 | 12.8 | 1.13 | 0.38 |
| SRR635193_1 | 1.47 | 0.6964 | 117.4 | 18.9 | 1.14 | 0.62 | 0.7023 | 408.1 | 33.9 | 5.60 | 1.97 | 0.6842 | 468.0 | 12.0 | 2.00 | 0.61 |
| MiSeq_1 | 0.87 | 0.2711 | 28.8 | 7.4 | 1.66 | 0.83 | 0.2584 | 63.7 | 10.4 | 2.70 | 0.97 | 0.2411 | 115.7 | 5.0 | 0.46 | 0.13 |
| SRR554369_1 | 0.17 | 0.4413 | 7.3 | 1.9 | 0.86 | 0.32 | 0.5094 | 25.8 | 2.7 | 1.74 | 0.20 | 0.4385 | 32.9 | 0.9 | 0.24 | 0.07 |

Compression ratios are in bits per base (bpb). “ctime” (resp. “dtime”) stands for compression (resp. decompression) time, and is expressed in seconds. “cmem” (resp. “dmem”) stands for peak compression (resp. decompression) memory usage and is expressed in gigabytes ( $=10^9$ bytes). Minicom failed to correctly decompress several datasets, which are marked with an asterisk. The name “MiSeq\_1” denotes the dataset MiSeq\_Ecoli\_MG1655\_110721\_PF\_1.

**Table 3.** Compression, time and memory usage in the order non-preserving regime on PE datasets

| dataset | size<br>[Gbs] | SPRING |  |  |  |  | Minicom |  |  |  |  | PgRC |  |  |  |  |
| --- | --- | --- | --- | --- | --- | --- | --- | --- | --- | --- | --- | --- | --- | --- | --- | --- |
|  |  | ratio | ctime | dtime | cmem | dmem | ratio | ctime | dtime | cmem | dmem | ratio | ctime | dtime | cmem | dmem |
| ERP001775 | 88.78 | 0.2305 | 14239.0 | 6677.0 | 46.62 | 6.25 | — | — | — | — | — | 0.2036 | 27010.0 | 1002.3 | 45.69 | 24.86 |
| ERR174310 | 41.93 | 0.3174 | 5357.0 | 1044.9 | 21.93 | 5.18 | 0.4810* | 23857.0 | 718.3 | 109.42 | 24.91 | 0.3071 | 18639.0 | 540.9 | 38.02 | 18.05 |
| ERR532393 | 7.15 | 0.5023 | 598.8 | 75.2 | 5.57 | 2.95 | 0.4713 | 847.1 | 84.6 | 19.04 | 4.29 | 0.4239 | 3291.3 | 61.5 | 10.23 | 2.74 |
| SRR065390 | 6.76 | 0.2677 | 667.4 | 58.0 | 3.78 | 1.84 | 0.2947* | 787.5 | 62.1 | 15.14 | 4.06 | 0.2091 | 1494.6 | 31.6 | 3.64 | 1.89 |
| SRR689233 | 2.95 | 0.2390 | 194.3 | 27.3 | 1.94 | 1.33 | 0.2570* | 196.1 | 30.1 | 7.71 | 1.97 | 0.2414 | 632.8 | 19.1 | 1.56 | 0.89 |
| SRR635193 | 2.94 | 0.3391 | 203.7 | 32.2 | 1.81 | 1.22 | 0.4187 | 778.1 | 31.3 | 9.56 | 1.97 | 0.3755 | 788.1 | 24.7 | 2.93 | 1.35 |
| MiSeq | 1.73 | 0.1994 | 99.3 | 14.2 | 1.86 | 1.50 | 0.1719 | 106.7 | 13.5 | 4.07 | 0.97 | 0.1516 | 296.1 | 8.5 | 0.79 | 0.34 |
| SRR554369 | 0.33 | 0.2221 | 12.9 | 3.5 | 0.94 | 0.57 | 0.2933 | 35.9 | 3.1 | 2.05 | 0.20 | 0.2207 | 53.0 | 1.5 | 0.32 | 0.11 |

Compression ratios are in bits per base (bpb). “ctime” (resp. “dtime”) stands for compression (resp. decompression) time, and is expressed in seconds. “cmem” (resp. “dmem”) stands for peak compression (resp. decompression) memory usage and is expressed in gigabytes ( $=10^9$ bytes). Minicom failed to compress one dataset, denoted with “—”, due to excessive memory usage, and failed to correctly decompress three other datasets, which are marked with an asterisk. The name “MiSeq” denotes the dataset MiSeq\_Ecoli\_MG1655\_110721\_PF.

**Table 4.** Compression, time and memory usage in the order preserving regime on PE datasets

| dataset | size<br>[Gbs] | SPRING |  |  |  |  | PgRC |  |  |  |  |
| --- | --- | --- | --- | --- | --- | --- | --- | --- | --- | --- | --- |
|  |  | ratio | ctime | dtime | cmem | dmem | ratio | ctime | dtime | cmem | dmem |
| ERP001775 | 88.78 | 0.3841 | 12582.0 | 3703.0 | 46.62 | 4.30 | 0.3409 | 32247.0 | 1437.4 | 49.33 | 26.21 |
| ERR174310 | 41.93 | 0.4633 | 4969.0 | 684.7 | 21.93 | 4.26 | 0.4395 | 21496.0 | 470.6 | 39.68 | 19.30 |
| ERR532393 | 7.15 | 0.6208 | 365.9 | 80.6 | 3.64 | 1.77 | 0.5381 | 3311.2 | 69.2 | 10.52 | 2.68 |
| SRR065390 | 6.76 | 0.3937 | 378.2 | 64.8 | 3.78 | 1.50 | 0.3236 | 1477.5 | 49.3 | 3.91 | 1.43 |
| SRR689233 | 2.95 | 0.3668 | 160.4 | 28.1 | 1.94 | 1.32 | 0.3566 | 650.0 | 26.4 | 1.74 | 0.70 |
| SRR635193 | 2.94 | 0.5617 | 176.7 | 35.1 | 1.80 | 1.01 | 0.5444 | 836.9 | 32.3 | 3.15 | 1.02 |
| MiSeq | 1.73 | 0.2738 | 66.2 | 14.5 | 1.86 | 1.48 | 0.2203 | 300.5 | 10.1 | 0.79 | 0.25 |
| SRR554369 | 0.33 | 0.3222 | 12.8 | 3.6 | 0.94 | 0.57 | 0.3133 | 53.4 | 2.5 | 0.33 | 0.09 |

Compression ratios are in bits per base (bpb). “ctime” (resp. “dtime”) stands for compression (resp. decompression) time, and is expressed in seconds. “cmem” (resp. “dmem”) stands for peak compression (resp. decompression) memory usage and is expressed in gigabytes ( $=10^9$ bytes). Minicom does not handle PE datasets in the order preserving regime. The name “MiSeq” denotes the dataset MiSeq\_Ecoli\_MG1655\_110721\_PF.

Let us first focus on the SE results. The average compression gain over the better of the two competitors (in the cases where Minicom failed only SPRING was used in the comparison) was over 11%. The gain over individual competitors was however greater: PgRC’s archive was on average by 17.7% smaller than SPRING’s and by 21.2% smaller than Minicom’s. The most striking improvement was achieved on SRR870667_1, where the PgRC archive is smaller than SPRING’s by 40.9% and than Minicom’s by 37.0%. The two cases where PgRC lost, only to SPRING, are ERR174310_1 and SRR554369_1, with 10.5% and 0.5% loss, respectively. In compression times, PgRC is much slower than its competitors, about 5 times than SPRING and more than 2.5 times than Minicom. This can be mostly explained by PgRC’s single-threaded implementation. Fortunately, things are not that bad in decompression time, where PgRC is, on average, by about 2.4 times faster than SPRING and 1.5 times faster than Minicom. PgRC is also more memory-frugal in compression (although not in decompression) than Minicom. SPRING needs least amounts of memory for the compression and the decompression, but from a practical point the most important values are peak memory usages for the largest datasets. To compress ERP001775_1, SPRING needs 23.5 GB of RAM, while PgRC spends 36.9 GB. In decompression, the corresponding peak memory usages are 5.3 GB and 16.4 GB, which should be acceptable for our compressor, assuming a mid-end desktop computer or a high-end laptop.

In the SE_ORD experiment, the general trend is similar, but the compression advantage of PgRC over its better competitor is smaller, 4.9% on average. As mentioned, also here PgRC produces the smallest archives, with two exceptions. On ERR174310_1 SPRING is the winner, while on SRR490976 Minicom takes the lead; in both cases the winner’s advantage over PgRC is about 4%. Concerning the compression speed, SPRING dominates again, being about 6 times faster than PgRC and about 2.5 times faster than Minicom. In decompression, SPRING is now usually faster than Minicom (with the exception of the largest dataset, ERP001775_1), but PgRC is again the fastest, about 1.6 (resp. 2.1) times faster than SPRING (resp. Minicom). Finally, the memory usage is almost unchanged for Minicom, with respect to the previously discussed SE mode, but grows somewhat for PgRC and, perhaps surprisingly, is reduced for SPRING (slightly in the compression phase and more significantly in the decompression). Yet, the peak memory usages for PgRC remain acceptable: 40.4 GB and 19.5 GB, respectively.

The last two experiments, with PE and PE_ORD modes, generally follow the pattern. PgRC wins in the compression ratio on 6 out of 8 datasets in the PE mode (while being defeated twice by SPRING) and on all 8 datasets in the PE_ORD mode. As Minicom fails in half of the cases of the former experiment (PE) and does not support PE_ORD, we now focus in a detailed comparison with SPRING only. The PgRC archives are on average by 8.2% smaller than SPRING’s in the PE mode and by 9.5% smaller in the PE_ORD mode. PgRC is on average 3.4 (resp. 5.1) times slower than SPRING in the compression in the PE (resp. PE_ORD) mode, but in the decompression PgRC is on average 2.3 (resp. 1.4) faster in the PE (resp. PE_ORD) mode. In compression and decompression memory SPRING is again more frugal, but, interestingly, for the largest dataset in the PE experiment PgRC spends slightly less memory at compression (45.69 GB vs 46.62 GB). In PE_ORD PgRC is worse in this aspect, with 49.33 GB of peak RAM usage, compared to 46.62 GB needed by SPRING.

We noticed that both SPRING and Minicom produce temporary disk files during the decompression, which stands in contrast to PgRC. To measure the impact of the storage medium, we compared the HDD used throughout our experiments against a fast SSD (with the sequential read and write performance up to 3400 MB/s and 1500 MB/s, respectively); the results of preliminary experiments are presented in Table 5. Only one dataset was used here, SRR065390, in two modes: PE and PE_ORD. As expected, PgRC is not much sensitive to the disk choice, although reading the input and writing the compressed output obviously benefit from the SSD also here. SPRING and Minicom perform more I/O operations and their compression and decompression times are reduced sometimes more than twice. We note in passing (cf. the first two rows in Table 5) that SPRING may produce slightly different archive sizes from run to run, presumably due to randomized hashing. The advantage of SPRING over PgRC in compression speed is thus even more pronounced in this scenario, while in decompression times now they are comparable.

**Table 5.**
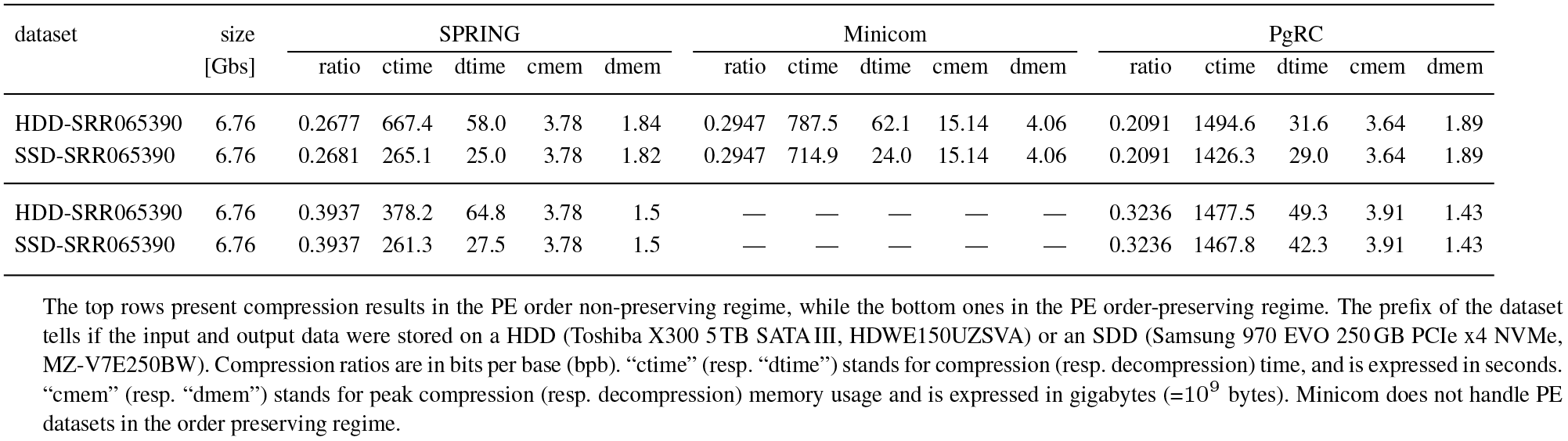
The impact of the disk (HDD vs SDD) on the compression, time and memory usage

To sum up the observations, our tool, PgRC, is significantly better in the compression ratio than its tested competitors, SPRING and Minicom, in all four compression modes, but it is also much inferior in compression speed, especially compared to SPRING. PgRC’s decompression speed and memory requirements are at least comparable to its competitors. Finally, we point out that we also compared PgRC to FQSqueezer, which failed to work on the largest datasets in the PE modes, due to excessive memory requirements. FQSqueezer, which is a multi-threaded tool, is comparable in compression speed to PgRC, but many times slower in decompression, due to the symmetric nature of the PPM compression approach. In compression ratio it wins over PgRC with the average archive smaller from 5.3% (PE_ORD) to 12.3% (PE), but its large computational requirements, especially on the receiver’s side, make it hardly practical, and this is why we present our experimental comparisons only in the Supplementary data.

## 4 Conclusions

We presented PgRC, a compression algorithm for the DNA stream of the FASTQ files, which achieves compression ratios significantly higher than its competitors, SPRING and Minicom. PgRC is also competitive in decompression speed and memory requirements, which makes it a viable choice for practical applications. The soft spot of our tool is compression speed, several times worse than of, e.g., SPRING, which is related to single-threaded implementation.

This weakness also suggests some lines for the future development. We are going to rewrite PgRC into a multi-threaded implementation. The proper choice of back-end compressors (currently, LZMA and PPMd from 7zip) is not obvious, concerning the time-space tradeoffs, but we believe we can make this final stage several times faster at preserved, or only slightly deteriorated, compression. PgRC makes use of several internal parameters, which were set experimentally. It might however be possible to tune them to a given dataset, with benefits in the overall time-compression relation. A crucial, but also often most time-consuming phase of PgRC compression is building the pseudogenomes. Orthogonal to a parallel architecture, a major boost is perhaps possible here due to algorithmic improvements. Finally, all those changes should be packaged into a full-fledged FASTQ compressor.

## Acknowledgements

This work was partially financed by the Smart Growth Operational Programme 2014-2020 project no POIR.04.01.02-00-0089/17-00 (the first author). The project is conducted in the Institute of Applied Computer Science at the Lodz University of Technology.

## Supplementary Material

### 1 Dataset URLs

Most of the dataset descriptions are taken from https://www.ebi.ac.uk.

ERP001775 (H. sapiens)

ftp://ftp.sra.ebi.ac.uk/vol1/fastq/ERR174/ERR174324/ERR174324_1.fastq.gz
ftp://ftp.sra.ebi.ac.uk/vol1/fastq/ERR174/ERR174325/ERR174325_1.fastq.gz
ftp://ftp.sra.ebi.ac.uk/vol1/fastq/ERR174/ERR174324/ERR174324_2.fastq.gz
ftp://ftp.sra.ebi.ac.uk/vol1/fastq/ERR174/ERR174325/ERR174325_2.fastq.gz

The first two files were concatenated and the last two files were concatenated to obtain a 28x coverage paired-end dataset.

ERR174310 (H. sapiens 1)

ftp://ftp.sra.ebi.ac.uk/vol1/fastq/ERR174/ERR174310/ERR174310_1.fastq.gz
ftp://ftp.sra.ebi.ac.uk/vol1/fastq/ERR174/ERR174310/ERR174310_2.fastq.gz

ERR532393 (Population Genomics of metagenome sequence)

ftp://ftp.sra.ebi.ac.uk/vol1/fastq/ERR532/ERR532393/ERR532393_1.fastq.gz
ftp://ftp.sra.ebi.ac.uk/vol1/fastq/ERR532/ERR532393/ERR532393_2.fastq.gz

MiSeq (E. coli)

ftp://webdata:webdata@ussd-ftp.illumina.com/Data/SequencingRuns/MG1655/MiSeq_Ecoli_MG1655_110721_PF_R1.fastq.gz
ftp://webdata:webdata@ussd-ftp.illumina.com/Data/SequencingRuns/MG1655/MiSeq_Ecoli_MG1655_110721_PF_R2.fastq.gz

SRR065390 (llumina Genome Analyzer II paired end sequencing)

ftp://ftp.sra.ebi.ac.uk/vol1/fastq/SRR065/SRR065390/SRR065390_1.fastq.gz
ftp://ftp.sra.ebi.ac.uk/vol1/fastq/SRR065/SRR065390/SRR065390_2.fastq.gz

SRR1294116 (Illumina HiSeq 2000 sequencing; GSM1395310: human ES cell line H9 (ucla h9); Homo sapiens; RNA-Seq)

ftp://ftp.sra.ebi.ac.uk/vol1/fastq/SRR129/006/SRR1294116/SRR1294116.fastq.gz

SRR445718 (Illumina HiSeq 2000 sequencing; GSM896803: Oocyte #1-C1; Homo sapiens; RNA-Seq)

ftp://ftp.sra.ebi.ac.uk/vol1/fastq/SRR445/SRR445718/SRR445718.fastq.gz

SRR445724 (Illumina HiSeq 2000 sequencing; GSM896809: 2-cell #1-C1; Homo sapiens; RNA-Seq)

ftp://ftp.sra.ebi.ac.uk/vol1/fastq/SRR445/SRR445724/SRR445724.fastq.gz

SRR445726 (Illumina HiSeq 2000 sequencing; GSM896811: 2-cell #2-C1; Homo sapiens; RNA-Seq)

ftp://ftp.sra.ebi.ac.uk/vol1/fastq/SRR445/SRR445726/SRR445726.fastq.gz

SRR490961 (Illumina HiSeq 2000 sequencing; GSM922146: 4-cell embryo#1-Cell#1; Homo sapiens; RNA-Seq)

ftp://ftp.sra.ebi.ac.uk/vol1/fastq/SRR490/SRR490961/SRR490961.fastq.gz

SRR490976 (Illumina HiSeq 2000 sequencing; GSM922161: 8-cell embryo#1-Cell#4; Homo sapiens; RNA-Seq)

ftp://ftp.sra.ebi.ac.uk/vol1/fastq/SRR490/SRR490976/SRR490976.fastq.gz

SRR554369 (P. aeruginosa)

ftp://ftp.sra.ebi.ac.uk/vol1/fastq/SRR554/SRR554369/SRR554369_1.fastq.gz
ftp://ftp.sra.ebi.ac.uk/vol1/fastq/SRR554/SRR554369/SRR554369_2.fastq.gz

SRR635193 (Illumina Genome Analyzer IIx paired end sequencing; pooled amnion from 5 term birth placentas)

ftp://ftp.sra.ebi.ac.uk/vol1/fastq/SRR635/SRR635193/SRR635193_1.fastq.gz
ftp://ftp.sra.ebi.ac.uk/vol1/fastq/SRR635/SRR635193/SRR635193_2.fastq.gz

SRR689233 (Illumina HiSeq 2000 paired end sequencing; GSM1080195: mouse oocyte 1; Mus musculus; RNA-Seq)

ftp://ftp.sra.ebi.ac.uk/vol1/fastq/SRR689/SRR689233/SRR689233_1.fastq.gz
ftp://ftp.sra.ebi.ac.uk/vol1/fastq/SRR689/SRR689233/SRR689233_2.fastq.gz

SRR870667 1 (T. cacao)

ftp://ftp.sra.ebi.ac.uk/vol1/fastq/SRR870/SRR870667/SRR8700667_1.fastq.gz

### 2 Tested programs

The following programs, with the set parameters, were used in our experiments, with their results presented either in the main paper or in Section 3 of the Supplementary data.

SPRING

(version from 2018-Dec-15, https://github.com/shubhamchandak94/Spring):

SE, compression of DNA stream:

~~~
./spring -c --no-ids --no-quality -i in.fastq -o comp.spring -t 12 -r
~~~

SE_ORD, compression of DNA stream:

~~~
./spring -c --no-ids --no-quality -i in.fastq -o comp.spring -t 12
~~~

PE, compression of DNA stream:

~~~
./spring -c --no-ids --no-quality -i in1.fastq in2.fastq -o comp.spring -t 12 -r
~~~

PE_ORD, compression of DNA stream:

~~~
./spring -c --no-ids --no-quality -i in1.fastq in2.fastq -o comp.spring -t 12
~~~

Minicom

(version from 2018-Dec-27, https://github.com/yuansliu/minicom):

SE, compression of DNA stream:

~~~
./minicom -r in.fastq -t 12
~~~

SE_ORD, compression of DNA stream:

~~~
./minicom -r in.fastq -t 12 -p
~~~

PE, compression of DNA stream:

~~~
./minicom -1 in1.fastq -2 in2.fastq -t 12
~~~

FQSqueezer v0.1

(version from 2019-Feb-24, https://github.com/refresh-bio/fqsqueezer):

SE, compression of DNA stream:

~~~
./fqs-0.1 e -im n -qm n -om s -s -t 12 -out comp.fqs in.fastq
~~~

SE_ORD, compression of DNA stream:

~~~
./fqs-0.1 e -im n -qm n -om o -s -t 12 -out comp.fqs in.fastq
~~~

PE, compression of DNA stream:

~~~
./fqs-0.1 e -im n -qm n -om s -p -t 12 -out comp.fqs in1.fastq in2.fastq
~~~

PE_ORD, compression of DNA stream:

~~~
./fqs-0.1 e -im n -qm n -om o -p -t 12 -out comp.fqs in1.fastq in2.fastq
~~~

PgRC v1.0

(https://github.com/kowallus/PgRC):

SE, compression of DNA stream:

~~~
./PgRC -i in.fastq -o comp.pgrc
~~~

SE_ORD, compression of DNA stream:

~~~
./PgRC -o -i in.fastq -o comp.pgrc
~~~

PE, compression of DNA stream:

~~~
./PgRC -i in1.fastq in2.fastq -o comp.pgrc
~~~

PE_ORD, compression of DNA stream:

~~~
./PgRC -o -i in.fastq -o comp.pgrc
~~~

### 3 Additional tables and figures

We start with comparing PgRC against FQSqueezer (Tables 1–4). Note that FQSqueezer is essentially a symmetric algorithm, with decompression faster than compression by usually only a few percent. On the other hand, we have to admit that FQSqueezer tends to be faster in compressing large datasets than PgRC, by 30–40 percent, and even 4 times faster on SRR870667 1 (SE and SE ORD modes). On smaller datasets, PgRC may win in compression speed, and its relatively best case also reveals an advantage by a factor of 4 (SRR689233, in the PE mode). The compression ratios that FQSqueezer attains are superior over the rest of the crowd, sometimes with about 20 percent (and more) advantage over the second contender (which is usually PgRC). In a few cases, however, PgRC takes the lead. FQSqueezer is rather demanding in memory usage, both in the compression and the decompression, and this is why we were not able to run it on a few largest PE datasets on our test machine with 128 GB of RAM. To sum up, we do not consider FQSqueezer a really practical alternative to best existing FASTQ compressors, yet its compression ratios may serve as a yardstick by which further progress in this domain should properly be measured.

Fig. 1 presents the impact of varying four parameters (*ov*, *s*, *M*, *p*) on the overall compression ratios and compression times, on three selected datasets. Let us remind that *ov* denotes the minimum allowed overlap in the read partitioning phase, and is expressed as a percentage of the read length (the read for which the found overlaps, both the left and the right one, are at least *ov* will be part of *PG_hq_*), *s* is the seed length in the read alignment phase, *M* is such that the maximum error level, i.e., the maximum allowed number of mismatches per aligned read, is ⌊*read len/M*⌋ (*M* = 6 for reads of length 100 implies that we allow up to 16 mismatches), and *p* is the minimum length of a reverse-complemented repeat. The default values of those parameters are 65, 38, 6 and 50, respectively.

Several observations can be made from this experiment. The default value of the parameter *ov* is perhaps satisfactory, but on two of the three datasets a slightly better compression (by around 0.5%) could be obtained for a smaller *ov* value, with a side effect of a few percent speedup.

**Figure 1:**
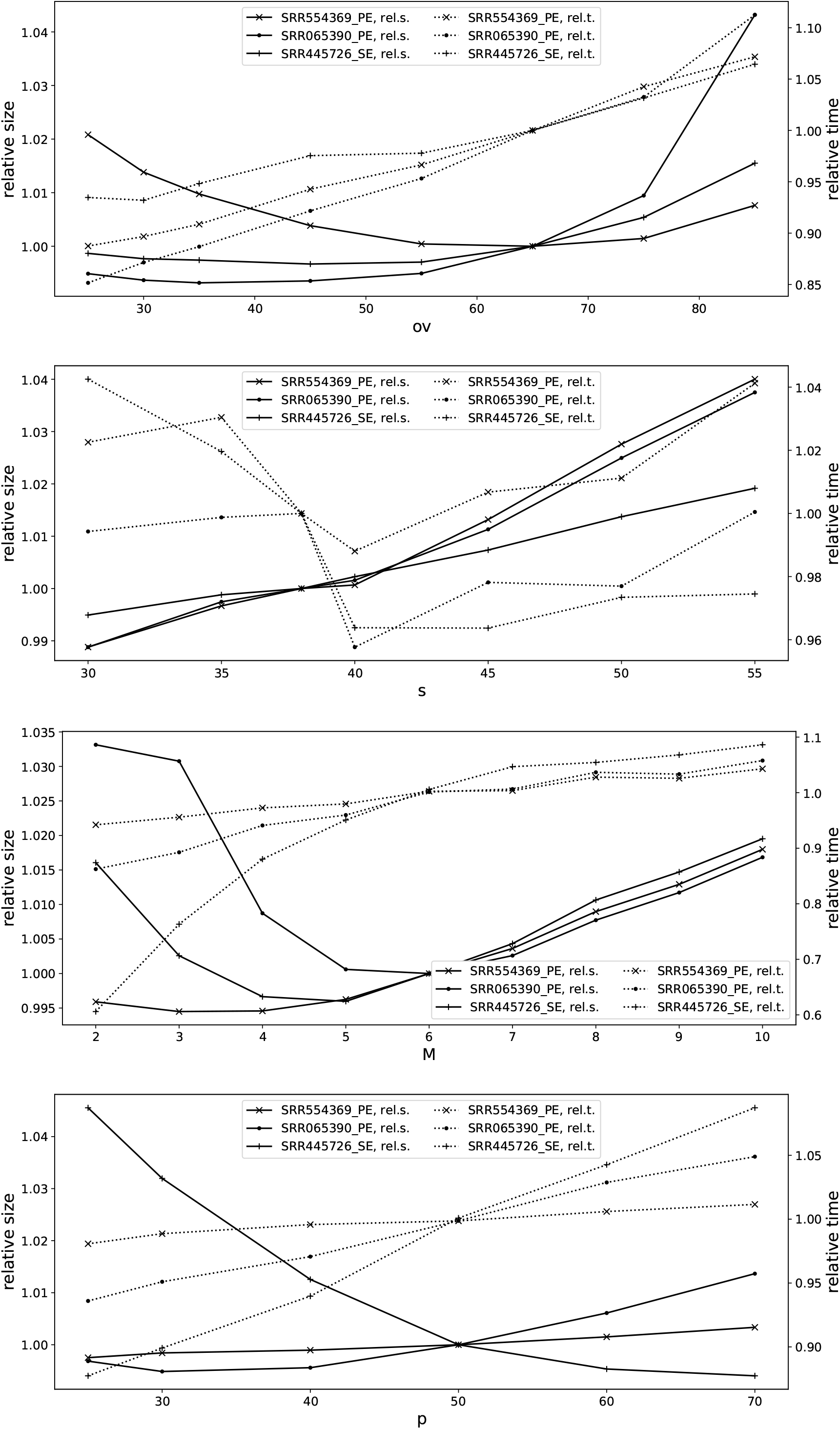
Relative compression ratios and times when one of the parameters is varied and the remaining three parameters keep their default value. The left (resp. right) Y axes are related to relative compressed sizes (resp. compression times). Three datasets were used: SRR554369_PE, SRR065390_PE, SRR445726_SE.

**Table 1:** Compression, time and memory usage in the order non-preserving regime on SE datasets. Compression ratios are in bits per base (bpb). “ctime” (resp. “dtime”) stands for compression (resp. decompression) time, and is expressed in seconds. “cmem” (resp. “dmem”) stands for peak compression (resp. decompression) memory usage and is expressed in gigabytes (=10^9^ bytes). The name “MiSeq_1” denotes the dataset MiSeq_Ecoli_MG1655_110721_PF_1.

| dataset | size<br>[Gbs] | Spring |  |  |  |  | FQSqueezer |  |  |  |  | PgRC |  |  |  |  |
| --- | --- | --- | --- | --- | --- | --- | --- | --- | --- | --- | --- | --- | --- | --- | --- | --- |
|  |  | ratio | ctime | dtime | cmem | dmem | ratio | ctime | dtime | cmem | dmem | ratio | ctime | dtime | cmem | dmem |
| ERP001775_1 | 44.39 | 0.2283 | 6283.0 | 1800.8 | 23.48 | 5.28 | 0.2480 | 15080.0 | 12833.0 | 97.65 | 95.19 | 0.2182 | 17410.0 | 423.4 | 36.92 | 16.43 |
| ERR174310_1 | 20.97 | 0.4096 | 2513.3 | 217.8 | 11.27 | 4.93 | 0.3928 | 10294.0 | 9282.0 | 93.82 | 92.74 | 0.4527 | 15197.0 | 135.5 | 21.17 | 12.35 |
| SRR870667_1 | 7.48 | 1.2924 | 1135.2 | 182.4 | 7.42 | 3.12 | 0.4997 | 3978.0 | 3628.0 | 69.38 | 69.06 | 0.7636 | 14151.0 | 79.2 | 19.95 | 13.76 |
| SRR445724 | 5.09 | 0.4328 | 685.8 | 70.0 | 3.47 | 2.52 | 0.2831 | 2038.5 | 1827.1 | 57.86 | 57.71 | 0.3444 | 2870.5 | 29.7 | 8.59 | 2.97 |
| SRR445726 | 4.92 | 0.3882 | 581.6 | 47.3 | 3.61 | 2.48 | 0.2608 | 1890.0 | 1690.7 | 57.15 | 57.00 | 0.3132 | 2472.8 | 27.3 | 7.57 | 2.59 |
| SRR490961 | 4.91 | 0.2183 | 467.3 | 40.9 | 2.92 | 2.15 | 0.1653 | 1489.1 | 1292.1 | 54.87 | 54.73 | 0.1841 | 1343.3 | 22.6 | 3.77 | 1.29 |
| SRR1294116 | 4.64 | 0.2763 | 304.9 | 41.9 | 3.19 | 2.32 | 0.2233 | 1732.9 | 1553.9 | 55.58 | 55.50 | 0.2415 | 1495.8 | 28.3 | 4.44 | 1.45 |
| ERR532393_1 | 3.58 | 0.4325 | 169.3 | 35.8 | 3.04 | 1.85 | 0.2968 | 1674.4 | 1533.3 | 55.22 | 55.16 | 0.3765 | 1749.7 | 21.9 | 6.18 | 1.36 |
| SRR065390_1 | 3.38 | 0.2275 | 175.3 | 28.6 | 2.63 | 1.24 | 0.1932 | 1245.2 | 1120.5 | 54.03 | 53.66 | 0.1877 | 773.1 | 17.9 | 2.09 | 0.79 |
| SRR490976 | 3.33 | 0.3851 | 407.2 | 32.4 | 2.84 | 2.14 | 0.3028 | 1606.3 | 1472.5 | 56.27 | 56.20 | 0.3411 | 1684.9 | 20.5 | 5.33 | 1.84 |
| SRR445718 | 3.29 | 0.3371 | 315.8 | 31.7 | 2.65 | 2.03 | 0.2529 | 1430.9 | 1303.0 | 55.37 | 55.26 | 0.2855 | 1365.7 | 18.6 | 4.02 | 1.33 |
| SRR689233_1 | 1.48 | 0.1934 | 66.0 | 14.1 | 1.61 | 0.80 | 0.1501 | 719.9 | 663.3 | 52.30 | 52.23 | 0.1700 | 330.9 | 6.2 | 0.90 | 0.33 |
| SRR635193_1 | 1.47 | 0.2671 | 108.2 | 16.2 | 1.30 | 0.81 | 0.2175 | 766.0 | 713.3 | 52.49 | 52.42 | 0.2599 | 438.1 | 7.9 | 1.78 | 0.53 |
| MiSeq_1 | 0.87 | 0.1253 | 36.6 | 7.3 | 1.66 | 0.84 | 0.0945 | 537.7 | 497.8 | 51.85 | 51.77 | 0.0947 | 110.1 | 2.2 | 0.39 | 0.11 |
| SRR554369_1 | 0.17 | 0.2397 | 7.2 | 1.8 | 0.86 | 0.32 | 0.2300 | 450.2 | 424.9 | 51.48 | 51.47 | 0.2409 | 32.2 | 0.7 | 0.23 | 0.06 |

**Table 2:** Compression, time and memory usage in the order preserving regime on SE datasets. Compression ratios are in bits per base (bpb). “ctime” (resp. “dtime”) stands for compression (resp. decompression) time, and is expressed in seconds. “cmem” (resp. “dmem”) stands for peak compression (resp. decompression) memory usage and is expressed in gigabytes (=10^9^ bytes). The name “MiSeq_1” denotes the dataset MiSeq_Ecoli_MG1655_110721_PF_1.

| dataset | size<br>[Gbs] | Spring |  |  |  |  | FQSqueezer |  |  |  |  | PgRC |  |  |  |  |
| --- | --- | --- | --- | --- | --- | --- | --- | --- | --- | --- | --- | --- | --- | --- | --- | --- |
|  |  | ratio | ctime | dtime | cmem | dmem | ratio | ctime | dtime | cmem | dmem | ratio | ctime | dtime | cmem | dmem |
| ERP001775_1 | 44.39 | 0.5275 | 4772.0 | 1426.8 | 23.48 | 4.30 | 0.5149 | 14108.0 | 13952.0 | 97.85 | 97.82 | 0.5039 | 22530.0 | 369.8 | 40.44 | 19.51 |
| ERR174310_1 | 20.97 | 0.6949 | 2509.0 | 248.5 | 11.27 | 4.43 | 0.6485 | 9390.0 | 9298.0 | 94.38 | 94.35 | 0.7236 | 17571.0 | 205.3 | 25.12 | 13.81 |
| SRR870667_1 | 7.48 | 1.4547 | 826.3 | 143.2 | 6.88 | 2.31 | 0.7221 | 3564.1 | 3548.7 | 69.60 | 69.59 | 1.0332 | 14604.0 | 102.7 | 20.48 | 14.24 |
| SRR445724 | 5.09 | 0.6574 | 606.4 | 64.6 | 2.90 | 1.12 | 0.5247 | 1817.8 | 1795.7 | 58.28 | 58.27 | 0.5984 | 2887.1 | 42.4 | 9.00 | 3.13 |
| SRR445726 | 4.92 | 0.6130 | 615.7 | 52.9 | 2.83 | 1.07 | 0.5029 | 1686.4 | 1667.2 | 57.61 | 57.59 | 0.5646 | 2505.7 | 39.4 | 7.97 | 2.73 |
| SRR490961 | 4.91 | 0.4572 | 423.0 | 47.2 | 2.83 | 0.83 | 0.4079 | 1320.9 | 1301.7 | 54.98 | 54.96 | 0.4336 | 1411.5 | 43.5 | 4.17 | 1.44 |
| SRR1294116 | 4.64 | 0.5122 | 213.3 | 58.1 | 2.70 | 0.90 | 0.4591 | 1504.5 | 1491.2 | 55.87 | 55.85 | 0.4829 | 1579.1 | 42.7 | 4.80 | 1.59 |
| ERR532393_1 | 3.58 | 0.6667 | 166.7 | 40.1 | 1.78 | 1.00 | 0.5325 | 1336.4 | 1328.9 | 55.26 | 55.24 | 0.6206 | 1774.5 | 30.2 | 6.47 | 1.47 |
| SRR065390_1 | 3.38 | 0.4776 | 175.4 | 32.7 | 2.63 | 0.83 | 0.4300 | 861.3 | 847.1 | 53.98 | 54.10 | 0.4278 | 805.2 | 19.4 | 2.47 | 0.89 |
| SRR490976 | 3.33 | 0.5979 | 435.3 | 36.6 | 2.23 | 1.11 | 0.5260 | 1209.7 | 1200.7 | 56.01 | 55.98 | 0.5879 | 1730.5 | 28.7 | 5.60 | 1.94 |
| SRR445718 | 3.29 | 0.5704 | 229.6 | 35.4 | 2.21 | 0.89 | 0.4871 | 1096.5 | 1086.7 | 55.23 | 55.22 | 0.5346 | 1392.4 | 32.8 | 4.28 | 1.43 |
| SRR689233_1 | 1.48 | 0.4489 | 66.1 | 14.5 | 1.61 | 0.74 | 0.4090 | 393.6 | 387.4 | 52.50 | 52.40 | 0.4303 | 347.5 | 12.8 | 1.13 | 0.38 |
| SRR635193_1 | 1.47 | 0.6964 | 117.4 | 18.9 | 1.14 | 0.62 | 0.6466 | 528.0 | 524.6 | 52.85 | 52.80 | 0.6842 | 468.0 | 12.0 | 2.00 | 0.61 |
| MiSeq_1 | 0.87 | 0.2711 | 28.8 | 7.4 | 1.66 | 0.83 | 0.2468 | 184.8 | 183.5 | 52.08 | 51.94 | 0.2411 | 115.7 | 5.0 | 0.46 | 0.13 |
| SRR554369_1 | 0.17 | 0.4413 | 7.3 | 1.9 | 0.86 | 0.32 | 0.4329 | 77.7 | 77.7 | 51.60 | 51.46 | 0.4385 | 32.9 | 0.9 | 0.24 | 0.07 |

Changing the parameter *s* is not easy: we can make the compression slightly better (again, a fraction of a percent), but also slightly slower, or make it faster by 2–4%, but with around 1% compression loss.

The choice for *M* is in two cases suboptimal and its smaller value (which means that more mismatches are allowed) could also make PgRC faster. The possible speedup is usually with 10%, but one dataset (SRR45726, the SE mode) reveals an interesting behavior: setting *M* to 2 would reduce the compression time by 40% at less than 2% compression loss; a tradeoff definitely worth considering.

Finally, playing around with *p* may give a slight improvement in the compression ratio (below 1%) with mixed impact on the compression time.

Fig 2 compares our ‘main’ strategy of dividing the reads into high quality and low quality ones (denoted as “pg generator” in the legend), with another, based on the FASTQ quality score stream (denoted as “quality stream”). The alternative approach is simple: the arithmetic mean of the characters encoding read’s quality scores is compared to a given threshold. This is equivalent to mapping the scores to probabilities and taking the geometric mean over them. Using different quality thresholds translates to a series of points varying with compression time and compression ratio. We can see that the solution based on FASTQ quality scores may be faster, in case of one of the shown datasets (SRR554369 PE) by even 30%, than our method of choice, but also produces archives larger by a few, or even more than 10 percent. For this reason, we believe that the chosen approach is superior. Note that manipulating with the *ov* parameter and thus obtaining the fraction of high quality reads different than the default may also speed things up a bit, but typically also with some (slight) penalty in compression ratio. Anyway, modifying this parameter may be an option for a fast compression mode.

Perhaps surprisingly, despite choosing a larger fraction of reads as high quality ones in the pg generation strategy, its resulting pseudogenome is typically much shorter. For example, for SRR554369 PE the pg generation strategy obtains the smallest archive with default settings and its pseudogenome size is 15.82 Mbases, while the smallest archive under the quality stream strategy is by 3.68% larger and its pseudogenome size is 18.62 Mb. The difference is more striking for SRR445726 SE, with 199.0 Mb in the former case and 7.01% and 495.7 Mb, respectively, in the latter case.

**Figure 2:**
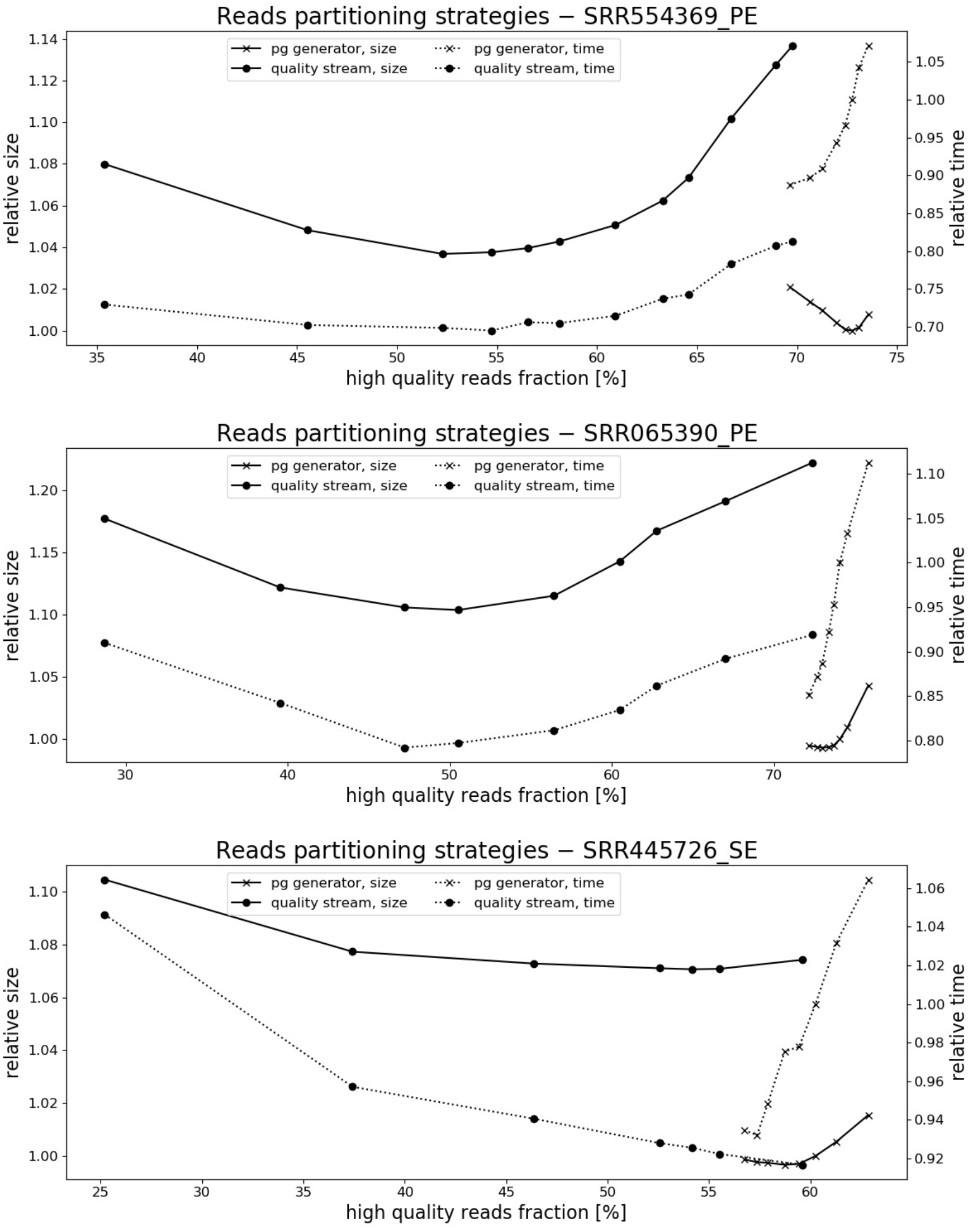
The impact of the method of dividing reads into high quality and low quality ones onto the resulting compression ratios and compression times. Three datasets were used: SRR554369_PE, SRR065390_PE, SRR445726_SE.

**Table 3:** Compression, time and memory usage in the order non-preserving regime on PE datasets. Compression ratios are in bits per base (bpb). “ctime” (resp. “dtime”) stands for compression (resp. decompression) time, and is expressed in seconds. “cmem” (resp. “dmem”) stands for peak compression (resp. decompression) memory usage and is expressed in gigabytes (=10^9^ bytes). FQSqueezer failed to compress two datasets, denoted with “—”, due to excessive memory usage. The name “MiSeq” denotes the dataset MiSeq_Ecoli_MG1655_110721_PF.

| dataset | size<br>[Gbs] | Spring |  |  |  |  | FQSqueezer |  |  |  |  | PgRC |  |  |  |  |
| --- | --- | --- | --- | --- | --- | --- | --- | --- | --- | --- | --- | --- | --- | --- | --- | --- |
|  |  | ratio | ctime | dtime | cmem | dmem | ratio | ctime | dtime | cmem | dmem | ratio | ctime | dtime | cmem | dmem |
| ERP001775 | 88.78 | 0.2305 | 14239.0 | 6677.0 | 46.62 | 6.25 | — | — | — | — | — | 0.2036 | 27010.0 | 1002.3 | 45.69 | 24.86 |
| ERR174310 | 41.93 | 0.3174 | 5357.0 | 1044.9 | 21.93 | 5.18 | — | — | — | — | — | 0.3071 | 18639.0 | 540.9 | 38.02 | 18.05 |
| ERR532393 | 7.15 | 0.5023 | 598.8 | 75.2 | 5.57 | 2.95 | 0.3535 | 2945.4 | 2641.4 | 73.38 | 73.18 | 0.4239 | 3291.3 | 61.5 | 10.23 | 2.74 |
| SRR065390 | 6.76 | 0.2677 | 667.4 | 58.0 | 3.78 | 1.84 | 0.2141 | 2254.8 | 2135.2 | 62.71 | 62.10 | 0.2091 | 1494.6 | 31.6 | 3.64 | 1.89 |
| SRR689233 | 2.95 | 0.2390 | 194.3 | 27.3 | 1.94 | 1.33 | 0.1859 | 2640.6 | 2508.7 | 57.26 | 57.18 | 0.2414 | 632.8 | 19.1 | 1.56 | 0.89 |
| SRR635193 | 2.94 | 0.3391 | 203.7 | 32.2 | 1.81 | 1.22 | 0.2988 | 1210.7 | 1096.1 | 61.24 | 61.22 | 0.3755 | 788.1 | 24.7 | 2.93 | 1.35 |
| MiSeq | 1.73 | 0.1994 | 99.3 | 14.2 | 1.86 | 1.50 | 0.1358 | 709.4 | 655.5 | 53.09 | 53.07 | 0.1516 | 296.1 | 8.5 | 0.79 | 0.34 |
| SRR554369 | 0.33 | 0.2221 | 12.9 | 3.5 | 0.94 | 0.57 | 0.2085 | 473.6 | 449.8 | 52.13 | 52.12 | 0.2207 | 53.0 | 1.5 | 0.32 | 0.11 |

**Table 4:** Compression, time and memory usage in the order preserving regime on PE datasets. Compression ratios are in bits per base (bpb). “ctime” (resp. “dtime”) stands for compression (resp. decompression) time, and is expressed in seconds. “cmem” (resp. “dmem”) stands for peak compression (resp. decompression) memory usage and is expressed in gigabytes (=10^9^ bytes). FQSqueezer failed to compress two datasets, denoted with “—”, due to excessive memory usage. The name “MiSeq” denotes the dataset MiSeq_Ecoli_MG1655_110721_PF.

| dataset | size<br>[Gbs] | Spring |  |  |  |  | FQSqueezer |  |  |  |  | PgRC |  |  |  |  |
| --- | --- | --- | --- | --- | --- | --- | --- | --- | --- | --- | --- | --- | --- | --- | --- | --- |
|  |  | ratio | ctime | dtime | cmem | dmem | ratio | ctime | dtime | cmem | dmem | ratio | ctime | dtime | cmem | dmem |
| ERP001775 | 88.78 | 0.3841 | 12582.0 | 3703.0 | 46.62 | 4.30 | — | — | — | — | — | 0.3409 | 32247.0 | 1437.4 | 49.33 | 26.21 |
| ERR174310 | 41.93 | 0.4633 | 4969.0 | 684.7 | 21.93 | 4.26 | — | — | — | — | — | 0.4395 | 21496.0 | 470.6 | 39.68 | 19.30 |
| ERR532393 | 7.15 | 0.6208 | 365.9 | 80.6 | 3.64 | 1.77 | 0.4698 | 2465.40 | 2447.90 | 72.93 | 72.78 | 0.5381 | 3311.2 | 69.2 | 10.52 | 2.68 |
| SRR065390 | 6.76 | 0.3937 | 378.2 | 64.8 | 3.78 | 1.50 | 0.3318 | 1865.70 | 1853.10 | 62.94 | 62.79 | 0.3236 | 1477.5 | 49.3 | 3.91 | 1.43 |
| SRR689233 | 2.95 | 0.3668 | 160.4 | 28.1 | 1.94 | 1.32 | 0.3151 | 2247.10 | 2230.80 | 57.50 | 57.33 | 0.3566 | 650.0 | 26.4 | 1.74 | 0.70 |
| SRR635193 | 2.94 | 0.5617 | 176.7 | 35.1 | 1.80 | 1.01 | 0.5136 | 961.40 | 952.10 | 61.26 | 61.10 | 0.5444 | 836.9 | 32.3 | 3.15 | 1.02 |
| MiSeq | 1.73 | 0.2738 | 66.2 | 14.5 | 1.86 | 1.48 | 0.2121 | 328.80 | 325.40 | 53.44 | 53.21 | 0.2203 | 300.5 | 10.1 | 0.79 | 0.25 |
| SRR554369 | 0.33 | 0.3222 | 12.8 | 3.6 | 0.94 | 0.57 | 0.3107 | 104.60 | 103.90 | 52.19 | 51.92 | 0.3133 | 53.4 | 2.5 | 0.33 | 0.09 |

Finally, we checked how much time of the total PgRC compression time is taken in particular phrases (Fig. 3) and what fraction of the whole archive particular components take (Fig. 4).

### 4 How PgRC handles the PE modes

In the modes PE and PE ORD, we need to remember the pairing information. In particular, in the PE mode we encode the location of the read stored further in the (conceptual) concatenation of the three pseudogenomes: *PG*_*hq*_, *PG*_*lq*_ and *PG*_*N*_, plus a flag telling if this read is taken from the file _1. In the PE ORD mode, however, we always encode the location of the read from the file _2.

The encoding algorithm for such snippets of data is based on two simple observations, concerning mostly pairs of high-quality reads, or more precisely, such pairs that both reads belong to *PG*_*hq*_.

- As the reads from _2 are reverse-complemented (at the start of their processing), many reads are in a close distance from their counterparts from the pair, a phenomenon analogous to the one for paired reads mapped onto a real genome.
- If, however, the distance between the two reads of a pair is large, then it happens quite often that such pairs of reads occur in adjacent positions, considering their relative order along the pseudogenome. Such a phenomenon usually corresponds to the case of having relatively long matching (directly or in the reverse-complemented manner) strings (possibly with few mismatches) in another region of the pseudogenome.

To address these redundancies, we proposed the following pairing information encoding algorithm, splitting the data into 7 streams. First we present them for the PE mode:

1. a binary flag telling if the distance between the two reads of the pair is small enough to fit a byte; the distance is expressed in the number of aligned reads between them (not in bases!),
2. if the answer is positive, the value (0 … 255) goes to Stream #2,
3. otherwise, a binary flag to tell if the difference between the current pair distance and the referential pair distance fits a byte (−128 … 127); we explain below what we mean by “referential pair”,
4. if the answer is positive, then mentioned difference, from −128 … 127, goes to the fourth stream,
5. otherwise, the current pair distance, represented in a 4-byte integer, goes to the fifth stream,
6. a binary flag only for a pair of reads close enough (flag “true” in Stream #1), telling if the first read of the pair, in the order of their location, comes from the file 1; this flag value is usually “true”,
7. a binary flag only for a pair of reads not close enough (flag “false” in Stream #1), telling if the first read of the pair, in the order of their location, comes from the file 1.

**Figure 3:**
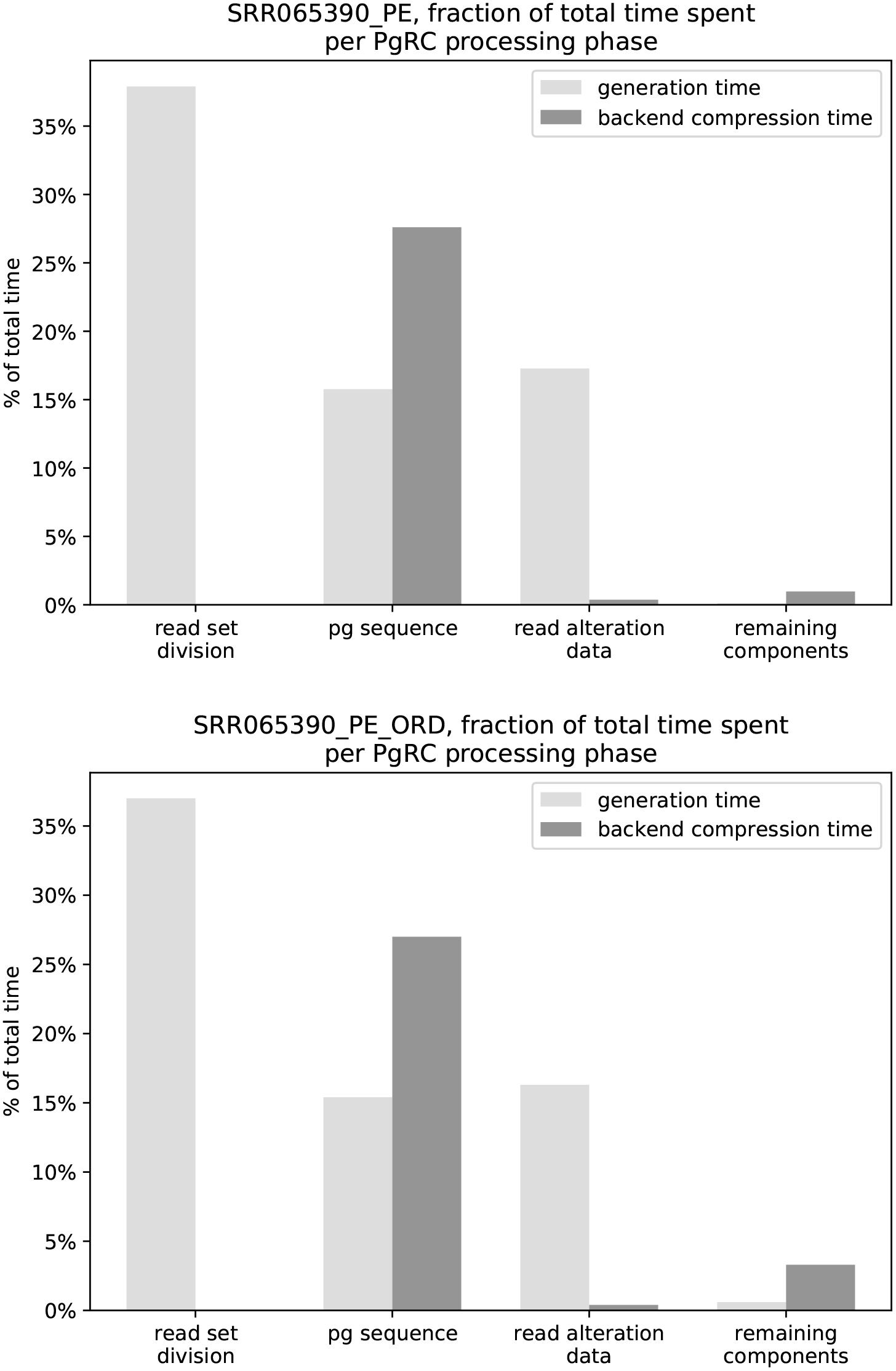
Fraction of the total time of PgRC’s compression process per phase. The top figure presents the results for SRR065390_PE (i.e., compressing SRR065390 in the order non-preserving PE mode) and the bottom one for SRR065390_PE_ORD. The bars are doubled: the left of each pair concerns producing “raw” components, and the correspond right one compressing it with LZMA or PPMd.

**Figure 4:**
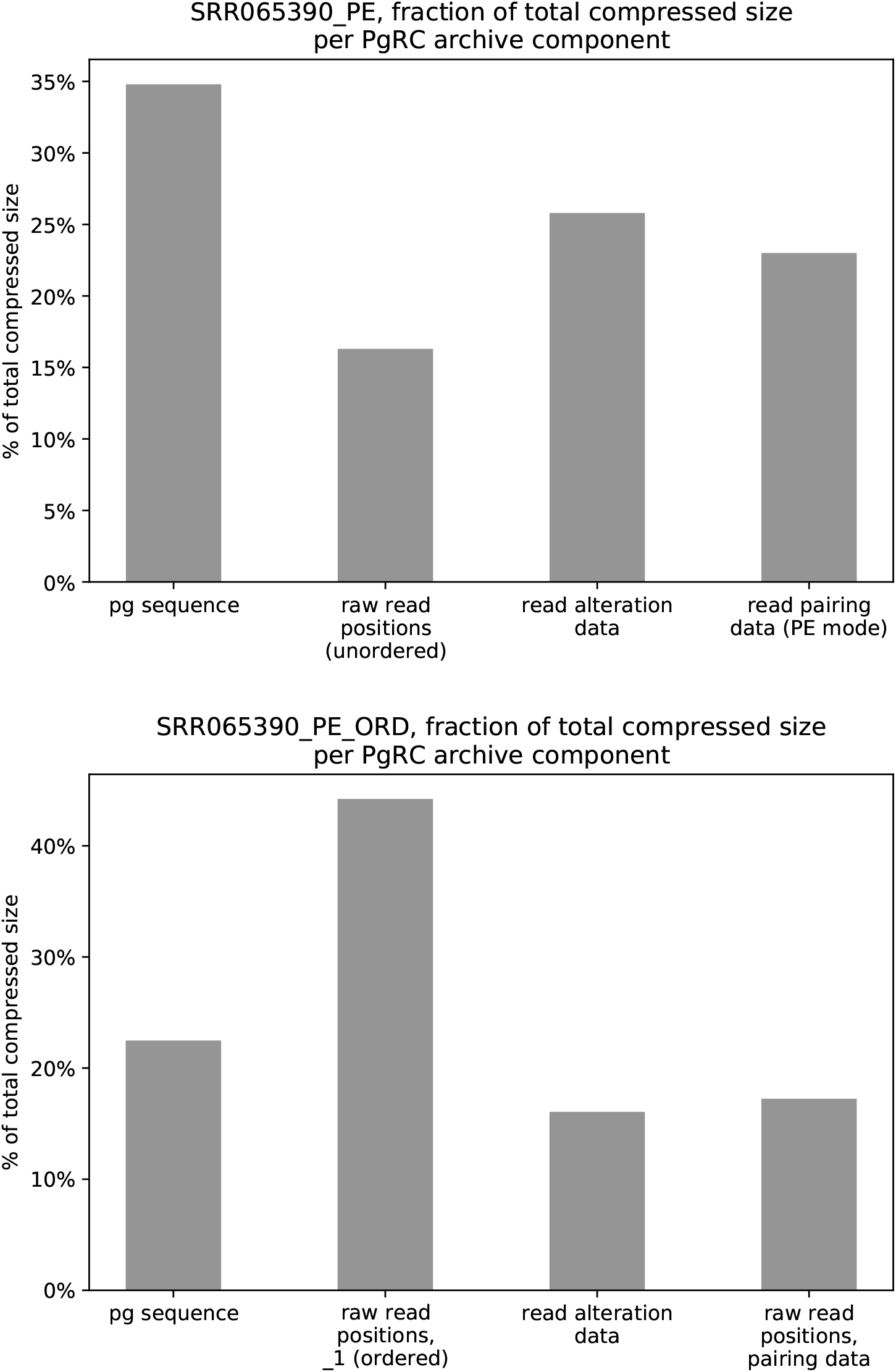
Fraction of the total size of PgRC’s archive per component. The top figure presents the results for SRR065390_PE (i.e., SRR065390 was compressed in the order non-preserving PE mode) and the bottom one for SRR065390_PE_ORD. Note that in the PE ORD mode the raw reads positions from the first file (_1), which are stored in the input FASTQ order, use 2.56 times more space (the second bar) than the extra information required to decode the _2 file pairing (the fourth bar).

The streams #4 and #5 are LZMA-compressed. The other streams are compressed with order-2 PPMd. The term “referential pair”, used in the description above, denotes either the currently last pair written to Stream #4, or the currently last pair written to Stream #5 provided that there has been at least two pairs written to Stream #5 after the last pair written to Stream #4.

## References

M. J. Bauer, A. J. Cox, and G. Rosone: Lightweight BWT Construction for Very Large String Collections. Proc. CPM 2011, pp. 219–231.

G. Benoit, C. Lemaitre, D. Lavenier, E. Drezen, T. Dayris, R. Uricaru, and G. Rizk: Reference-free compression of high throughput sequencing data with a probabilistic de Bruijn graph. BMC Bioinformatics, 16:288 2015.

J. K. Bonfield and M. V. Mahoney: Compression of FASTQ and SAM format sequencing data. PloS ONE, 8(3) 2013, pages e59190.

S. Chandak, K. Tatwawadi, I. Ochoa, M. Hernaez, and T. Weissman: SPRING: A next-generation compressor for FASTQ data. Bioinformatics, bty1015, https://doi.org/10.1093/bioinformatics/bty1015.

S. Chandak, K. Tatwawadi, and T. Weissman: Compression of genomic sequencing reads via hash-based reordering: algorithm and analysis. Bioinformatics, 34(4) 2018, pp. 558–567.

A. J. Cox, M. J. Bauer, T. Jakobi, and G. Rosone: Large-scale compression of genomic sequence databases with the Burrows–Wheeler transform. Bioinformatics, 28(11) 2012, pp. 1415–1419.

S. Deorowicz and S. Grabowski: Compression of DNA sequence reads in FASTQ format. Bioinformatics, 27(6) 2011, pp. 860–862.

S. Deorowicz: FQSqueezer: *k*-mer-based compression of sequencing data. bioRxiv, 10.1101/559807, 2019.

H. Y. Fritz, R. Leinonen, G. Cochrane, and E. Birney: Efficient storage of high throughput DNA sequencing data using reference-based compression. Genome Research, 21(5) 2011, pp. 734–740.

A. A. Ginart, J. Hui, K. Zhu, I. Numanagić, T. A. Courtade, S. C. Sahinalp, and D. N. Tse: Optimal compressed representation of high throughput sequence data via light assembly. Nature Communications, 9:566 2018.

S. Grabowski, S. Deorowicz, and Ł. Roguski: Disk-based compression of data from genome sequencing. Bioinformatics, 31(9) 2015, pp. 1389–1395.

S. Grabowski, W. Bieniecki: copMEM: finding maximal exact matches via sampling both genomes. Bioinformatics, 35(4) 2019, pp. 677–678.

F. Hach, I. Numanagić, C. Alkan, and S. C. Sahinalp: SCALCE: boosting sequence compression algorithms using locally consistent encoding. Bioinformatics, 28(23), pp. 3051–3057.

M. Howison: High-Throughput Compression of FASTQ Data with SeqDB. IEEE/ACM Transactions on Computational Biology and Bioinformatics, 10(1) 2013, pp. 213–218.

D. C. Jones, W. L. Ruzzo, X. Peng, M. G. Katze: Compression of next-generation sequencing reads aided by highly efficient de novo assembly. Nucleic Acids Research, 40(22), 2012, e171.

C. Kingsford and R. Patro: Reference-based compression of short-read sequences using path encoding. Bioinformatics, 31(12), 2015, pp. 1920–1928.

T. Kowalski, S. Grabowski, and S. Deorowicz: Indexing Arbitrary-Length *k*-Mers in Sequencing Reads. PLoS ONE, 10(7):e0133198 2015.

Y. Liu, Z. Yu, M. E. Dinger, and J. Li: Index suffix-prefix overlaps by (*w, k*)-minimizer to generate long contigs for reads compression. Bioinformatics, bty936, https://doi.org/10.1093/bioinformatics/bty936.

D. Maier and J. Storer: A note on the complexity of the superstring problem. Technical Report of Department of Electrical Engineering and Computer Science 233, Princeton University.

I. Ochoa, M. Hernaez, R. Goldfeder, T. Weissman, and E. Ashley: Effect of lossy compression of quality scores on variant calling. Briefings in Bioinformatics, 18(2), 2017, pp. 183–194.

R. Patro and C. Kingsford: Data-dependent bucketing improves reference-free compression of sequencing reads. Bioinformatics, 31(17), 2015, pp. 2770–2777.

M. Roberts, W. Hayes, B. R. Hunt, S. M. Mount, and J. A. Yorke: Reducing storage requirements for biological sequence comparison. Bioinformatics, 20(18) 2004, pp. 3363–3369.

Ł. Roguski and S. Deorowicz: DSRC 2—Industry-oriented compression of FASTQ files. Bioinformatics, 30(15) 2014, pp. 2213–2215.

Ł. Roguski, I. Ochoa, M. Hernaez, and S. Deorowicz: FaStore: a space-saving solution for raw sequencing data. Bioinformatics, 34(16) 2018, pp. 2748–2756.

H. Sarkar and R. Patro: Quark enables semi-reference-based compression of RNA-seq data. Bioinformatics, 33(21) 2017, pp. 3380–3386.

W. Tembe, J. Lowey, and E. Suh: G-SQZ: compact encoding of genomic sequence and quality data. Bioinformatics, 26(17) 2010, pp. 2192–2194.

V. Yanovsky: ReCoil—an algorithm for compression of extremely large datasets of DNA data. Algorithms for Molecular Biology, 6:23 2011.

Y. Zhang, L. Li, Y. Yang, X. Yang, S. He, and Z. Zhu: Light-weight reference-based compression of FASTQ data. BMC Bioinformatics, 16(1):188 2015.

